# Progressive maturation of the root apical meristem in *Arabidopsis thaliana* lateral roots

**DOI:** 10.1101/2022.02.18.481036

**Authors:** Béatrice Berthet, Lotte Bald, Marion Louveaux, Alexis Maizel

**Author notes:** **Emails:** Béatrice Berthet, Lotte Bald, Marion Louveaux, Alexis Maizel.

## Abstract

Meristems are stem cells niches that support the formation of all plant organs and are either set during embryogenesis and maintained throughout the plant life or specified de novo, post-embryonically. The embryo-derived root apical meristem is organized around a group of infrequently dividing cells, the quiescent centre, that maintains the stem cells, organizes growth along two axes and owing to its resistance to 3ic stress can replace damaged stem cells. In most cases, lateral roots post-embryonically branch off the primary and establish a new root meristem which organization is identical to the primary root one. The cellular and molecular processes underpinning the emergence of new stem cell niches are not well known. Here, we characterize the de novo establishment of the root apical meristem in lateral roots. While the position of the new stem cell niche is set early during morphogenesis, its cellular layout, unique gene expression profile and mitotic quiescence are only acquired after emergence concomitant to the establishment of two diverging growth axis. Our results show that the intertwined attributes of the mature root stem cell niche are progressively acquired during lateral root formation, and support a model in which the position of the stem cell niche emerges from the establishment of diverging growth axis.

**Highlight:** Analyze of the ontogeny of the quiescent center during lateral root ontogeny reveal its late formation and supports that its emergence results from the establishment of two diverging growth axis.

## Introduction

Above and below ground axial growth of plants is supported by primary meristems that are established during embryogenesis activated upon germination (Greb and Lohmann, 2016; ten Hove et al., 2015). Post embryonically, secondary meristems are established and allow branching and radial growth (Greb and Lohmann, 2016). These secondary meristems have a different ontogeny from their embryonic counterparts, but their organization and function is very comparable to the primary ones. They are thus very useful for elucidating the mostly unknown cellular and molecular processes governing the generation of new meristems (Greb and Lohmann, 2016). Lateral root (LR) branching from the primary or other root is an example of an organ which growth is supported by a secondary meristem (Motte *et al*., 2019). In *Arabidopsis thaliana (Arabidopsis)*, LR initiate deep within the primary root, where a few cells of the pericycle adjacent to the xylem radially expand and divide asymmetrically in response to local accumulation of the phytohormone auxin. These cells further grow, alternate rounds of anticlinal and periclinal divisions and form a dome shaped LR primordium (LRP) with a regular cellular layout which allows classification of its development in seven developmental stages (stage I to stage VII) (Malamy and Benfey, 1997; von Wangenheim *et al*., 2016). Ultimately, the new LR forms a new meristem with a cellular and functional organization quasi identical to the one of the primary root meristem (Tian et al., 2014b).

Meristems contain stem cells, also called initials, which activity is maintained by a group of rarely or non-dividing cells forming the organising centre in the shoot and the quiescent centre (QC) in the root (Dubrovsky and Ivanov, 2021). In *Arabidopsis*, the cellular organization of the root apical meristem (RAM) is simple and stereotyped with the central stem cells surrounding the QC. The stem cells, also called initials, provide, through asymmetric divisions, the progenitors for the root tissues and the root cap (Dolan *et al*., 1993). The concept of QC was formulated by F. A. L. Clowes (Clowes, 1956) and originally referred to both the organizer and the initials (Dubrovsky and Ivanov, 2021). A more restrictive definition of the QC, that only refers to the four to six apically located very slowly dividing cells was later adopted in *Arabidopsis* (Dolan *et al*., 1993; Dubrovsky and Ivanov, 2021). Here, we will use this latter definition for the QC. In addition to its low mitotic activity (Clowes, 1954; Clowes, 1956), the *Arabidopsis* QC is defined by additional features. First, it acts as an organiser that maintains the stem cell identity of cells in its direct contact (van den Berg et al., 1995; van den Berg et al., 1997). Second, it is characterized by abundance of auxin, an oxidizing environment and enrichment for expression of many genes and markers (Brumos et al., 2018; Haecker et al., 2004; Jiang et al., 2003; Nawy et al., 2005; Petersson et al., 2009; Sabatini et al., 1999). Third, the QC serves as a reservoir of cells to allow the meristem to recover from wounding or severe genotoxic damage by replacing compromised or missing cells (Clowes, 1959; Efroni et al., 2016; Fulcher and Sablowski, 2009; Tsugeki and Fedoroff, 1999). Thus, the QC defines through its specific central location a microenvironment, akin to a niche, that maintains the cellular organization of the meristem (Dubrovsky and Ivanov, 2021). The embryonic origin of the QC in *Arabidopsis* can be traced back to the late globular stage and the division of the hypophysis (Dolan *et al*., 1993; Scheres *et al*., 1994; ten Hove *et al*., 2015). Studies in maize indicate that all cells of the embryonic root proliferate and that quiescence of the QC is only established late during embryogenesis (Clowes, 1978; Dubrovsky and Ivanov, 2021).

In LR, seminal works hint at a late formation of the QC in the water cabbage *Pistia*, the water hyacinth *Eichhornia*, bean (*Vicia faba)*, mallow *(Malva sylvestris)* and Maize (Byrne, 1973; Clowes, 1958; Clowes, 1978; Macleod, 1977). In Arabidopsis, it was proposed that the formation of the RAM in lateral root only occurs after an initial early morphogenesis phase (up to stage IV/V) (Laskowski *et al*., 1995). Similarly, RAM formation in adventitious roots has been reported to take place late (Della Rovere *et al*., 2013). Using reporters for QC markers, Goh et al. traced back the origin of QC cells to the stage II LRP cells specifically expressing the *SCARECROW* (*SCR*) transcription factor. Disrupting *SCR* function perturbed the formation of these QC precursor cells (Goh *et al*., 2016). A recent analysis of LR formation at single cell resolution (Serrano-Ron *et al*., 2021) revealed that although expression of QC markers debuts early during LR formation they are distributed amongst different cell populations and a distinct coherent population only emerges at later stages.

Thus, whereas the cellular origin of the QC and the expression of QC markers are set early during RAM formation in LR, at which stage a fully functional stem cell niche is established remains elusive. Here, we investigate in detail the establishment of the lateral root stem cell niche. In particular we focus on the ontogeny of the QC, considering when each of its features are established: position, slow proliferation, expression of specific markers, and maintenance of the stem cells. We use a combination of tissue identity reporters, live imaging, lineage tracing and cell proliferation assay to conclude that the maturation of the stem cell niche is a progressive process during LR development with acquisition of QC position, molecular identity, quiescence and function being temporally uncoupled. While the QC position, and its role in proximal (shootward) growth are set early during LR formation, its molecular identity, cellular organization and quiescence are only fixed after LR emergence.

## Results

### Classification of lateral root post-emergence

Prior to emergence, LR development follows a nomenclature based on standard morphological stages (Stage I to VII) (Malamy and Benfey, 1997). For the post emergence phases, six types of lateral roots are defined based on their length and angle to the primary root (Kiss *et al*., 2002). Here, we adapted this latter nomenclature to microscopy image, measuring LR from the junction between the vasculature of the LR and the primary root to the tip of the LR. While we refer to type 1 as pre-emerged LR, type 2 LR have a maximal length of 250 µm; type 3 a length of up to 1 mm, type 4 up to 2 mm and type 5, more than 2 mm (Fig. 1A).

**Fig. 1.**
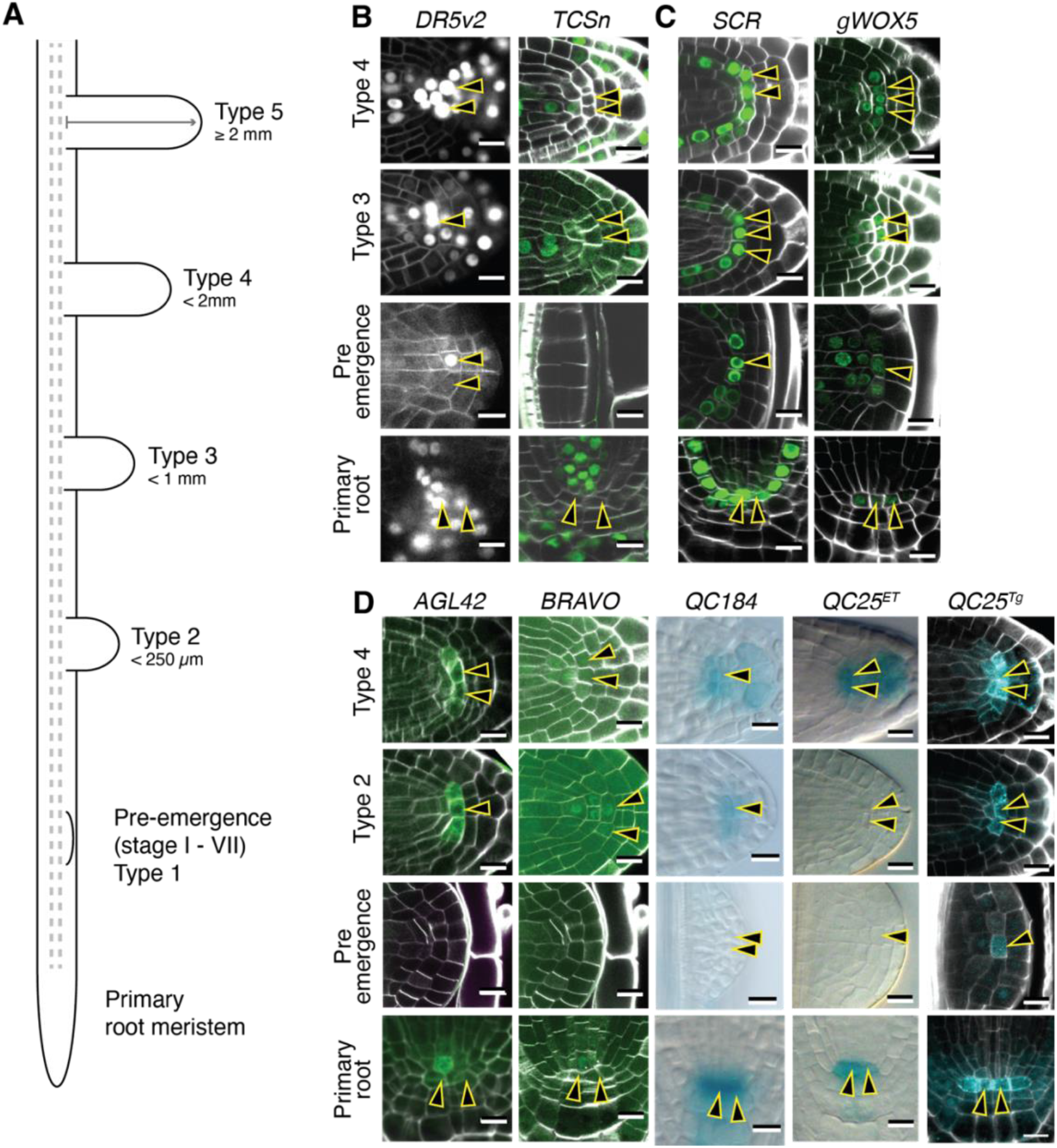
Position and identity of the stem cell niche during lateral root development. (A) Classification of emerged LR based on their length measured from the xylem strands of the primary root to the tip of the lateral root. (B-D) Micrographs of mature primary and developing lateral roots meristems expressing reporters for (B) auxin (*DR5v2*) or cytokinin (*TCSn*) signalling, (C) for the niche position (*gWOX5* or *SCR*), (D) for QC specific markers (*AGL4, BRAVO, QC184, QC25*^*ET*^, *QC25*^*Tg*^). *SCR* is a transcriptional reporter (*pSCR::nGFP*), *gWOX5* is a YFP-tagged genomic clone (*WOX5-YFP), AGL42* and *BRAVO* are translational reporters. *QC184* and *QC25*^*ET*^ are *GUS* enhancer trap lines expressing while in *QC25*^*TG*^ erCFP is expressed from the cis regulatory sequence located upstream of the *QC25*^*ET*^ insertion. Images are confocal sections of fluorescent reporters counterstained with Calcofluor white except for *QC184* and *QC25*^*ET*^ that are widefield images of GUS reporters. The apically located cells are indicated with arrowheads. Scale bars: 10μm.

### The position of the lateral root stem cell niche is defined before emergence, its identity after

In the primary root, the stem cell niche is positioned by complementary patterns of auxin and cytokinin accumulation (Bishopp et al., 2011; Müller and Sheen, 2008; Salvi et al., 2020). There, auxin abundance and auxin signalling are maximal in the QC and the columella cells (Sabatini *et al*., 1999) while cytokinins signalling is maximal in the root cap, the pro-vasculature above the QC and the distal region of the differentiation zone (Bielach *et al*., 2012; Marhavý *et al*., 2014; Montesinos *et al*., 2020). Auxin and cytokinin tightly regulate each phase of LR development (Nenadić and Vermeer, 2021). We set to see if in the developing LR the position of the new stem cell niche is defined by the same complementary pattern. We visualized the output of auxin and cytokinin signalling during LR development using the *pDR5v2::NLS-3xVenus* (*DR5v2::Venus*, (Liao *et al*., 2015) and *TCSn::2xVenus-NLS* (*TCSn::Venus*, (Zürcher *et al*., 2013) reporters (Fig. 1B). Before emergence, *DR5v2::Venus* signal is present throughout the LRP up to stage IV and after becomes restricted to the tip, as previously observed (Benková *et al*., 2003). Post emergence, *DR5v2::Venus* expression is similar to the pattern of the primary root, with highest signal at the apex of the meristem and the columella. *TCSn::Venus* expression was not detected in the LRP before emergence. This result confirms the previous observations of low level of cytokinin response in the LRP before emergence made with the first generation TCS reporter (Bielach et al., 2012; Chang et al., 2015; Müller and Sheen, 2008). In type 2 emerged LR, *TCSn::Venus* expression is first detected in the LR stele, then in the columella. In longer LR, *TCSn::Venus* expression is visible in the outermost layers of the LR columella, an expression pattern reminiscent of the primary root one. Thus while auxin signalling pattern marks the position of the LR stem cell niche from stage IV onward, the complementary cytokinin signalling pattern is only set post emergence.

In the primary root, the expression of *SCARECROW* (*SCR*) and *WOX5* overlap at the quiescent centre (Shimotohno *et al*., 2018). In LRP, an analysis of the expression pattern of *SCR* and *WOX5* using transcriptional reporters concluded that the LR stem cell niche is established by stage IV, with *WOX5* specifically expressed in the apically located cells of the second outermost layer at stage IV/V (Goh *et al*., 2016). We imaged a *SCR* transcriptional reporter (*pSCR::nGFP*) and confirmed it was first detected in the outer layer of stage II LRP and in later stages formed the typical arch marking the stem cell niche and endodermis layer, a profile maintained post emergence (Fig. 1C). We also imaged a YFP-tagged *WOX5* genomic clone (*WOX5-YFP*, (Pi *et al*., 2015)) and, unlike what was previously reported (Goh *et al*., 2016), we readily observed *WOX5-YFP* expression in all cells of stage I-III primordia. From stage IV onward, *WOX5-YFP* was restricted to the centre of the primordium (Supplemental Fig. S1 and Fig. 1C). Post emergence, *WOX5-YFP* expression is progressively restricted to three to five cells at the apex of type 4 emerged LR (Fig. 1C). Together, these results suggest that the position of the LR stem cell niche, defined by the overlap of an auxin maximum and expression of *WOX5* and *SCR* is defined prior stage IV of LR development, but is only established with cellular precision after emergence, suggesting its identity is only set then.

To verify that a specific niche identity comparable to the primary root is only acquired post-emergence, we monitored the expression of several markers specific of the QC in the primary root. We used two enhancer traps, QC184 and QC25^ET^ (Sabatini *et al*., 2003) and two transcription factors, BRAVO (Vilarrasa-Blasi *et al*., 2014) and AGL42 (Nawy *et al*., 2005). Before LR emergence, none of these markers were detected (Fig. 1D). They became visible in type 2 (AGL42, BRAVO, QC184) or type 3 (QC25^ET^) LR (Fig. 1D). The expression of QC25 was previously reported in pre-emergence LR, in two apically located cells from stage IV onward (Goh *et al*., 2016), much earlier than seen here. However the QC25 reporter used in that study is a transgenic line (QC25^Tg^, (ten Hove *et al*., 2010)) which was constructed by isolation of the sequence flanking the insertion site of the QC25^ET^ enhancer trap used (QC25^ET^) here. Whereas in the primary root both versions mark identically the QC, in the developing LR, QC25^ET^ only marks the apically located cells of the niche post-emergence, suggesting, that complex genomic regulation might be at play. Together, these results indicate that the position of the LR root apical meristem is defined before emergence whereas it only expresses specific QC markers once the LR have emerged.

### The cellular layout of the LR stem cell niche is set post-emergence

We then asked when during LR ontogeny, the typical cellular organization of the meristem is established. The QC of the primary root consists of six to twelve apically located cells arranged in a monolayered disk (Dolan *et al*., 1993; Lu *et al*., 2021). On sections, two to four apically located cells are typically visible and located between the two cortex-endodermis initials (Fig. 2A). To determine at which stage of LR development this organising centre is first visible and how many cells it has, we imaged by confocal microscopy Calcofluor White stained (Ursache *et al*., 2018) LRP of 10-day-old *Arabidopsis*. Prior to emergence, apically located cells between the two cortex-endodermis initials could only be unambiguously observed from stage V onward (Fig. 2A). Until emergence, their number was variable, with two cells visible only in 40% of the LRP sections (n=105, Fig. 2B). Post emergence, this variability progressively diminished with most type 4 LR having two cells visible, like in the primary root (Fig. 2B). We confirmed this progressive maturation of the LR stem cell niche by counting the number of QC25^Tg^ forming a disk on transverse section (Fig. 2C, D). While pre-emergence, less than 5 cells were typically observed, this number increased as the LR mature post-emergence to reach the values seen in the primary root meristem (Fig. 2D). Together, these data show the typical cellular layout of the root meristem is only fixed after emergence and indicates that the cellular organization of the LR organising centre progressively matures. The primary root meristem has been shown to progressively acquire its organization post-germination (Salvi *et al*., 2020). We thus asked whether the maturation of the LR stem cell niche post-emergence follows a similar path to the primary root meristem post-germination. We counted the number of apically located cells in the primary root meristem during germination and observed that on average two cells were already visible on section at 0, 24 or 48h post imbibition (Fig. S2). Thus the establishment of the cellular layout of the primary and lateral root stem cell niches follow different path.

**Fig. 2.**
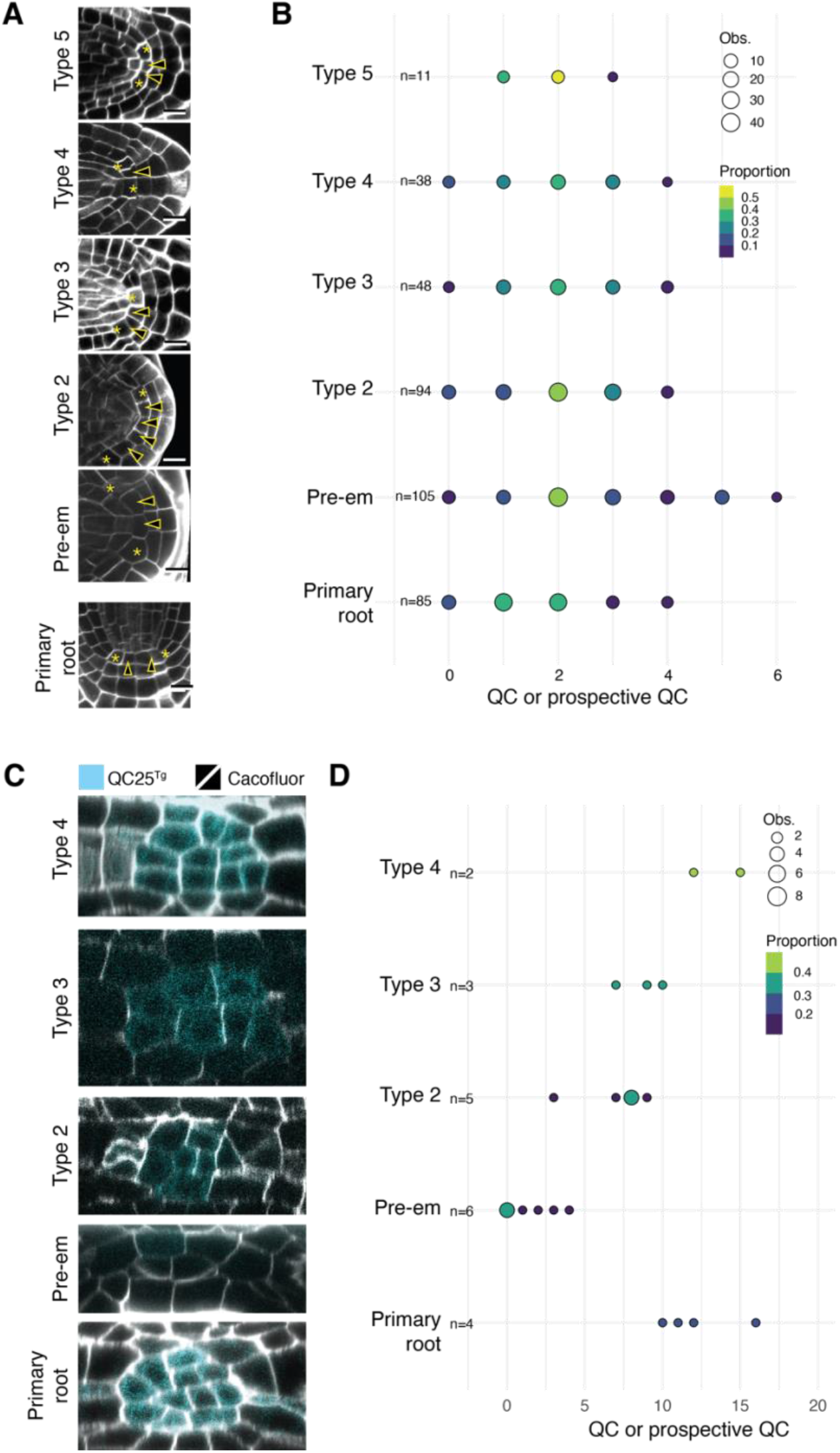
Cellular layout of developing apical meristem of the lateral root. (A) Confocal sections of the developing root apical meristem and each class of LR counterstained with Calcofluor white in 10-day-old Arabidopsis. The QC or apically located prospective QC cells (arrowheads) and the cortex-endodermal initial (stars) are indicated. Scale bars: 10μm. (B) Number of QC or apically located prospective QC cells on sections for the different classes of roots. (C) Cross-sections of 10-day-old *QC25*^*Tg*^ seedlings counterstained with Calcofluor white. (D) Total number of QC cells on sections for the different classes of roots. In B & D, the size of the circle is proportional to the number of observations (Obs.), and the colour represents the relative proportion of observations within a given class. The total number of observations per given classes is indicated (n).

### The LR stem cell niche becomes quiescent shortly after emergence

A distinctive feature of the primary root meristem is the low mitotic activity of the QC cells. We thus asked when cells become quiescent during LR formation. For this we quantified the proportion of proliferating apically located cells using the G2-S phase label 5-ethynyl-2’-deoxyuridine (EdU) during LR ontogeny, before and after emergence. Plants expressing the *pSCR::nGFP* reporter were incubated with EdU for 8h, fixed, and LR were imaged. We quantified the fraction of *SCR* positive cells in between the two cortex-endodermis initials (apically located cells) that are EdU positive (Fig. 3A). Before emergence, >75% of apically located cells of stage IV to VI primordia incorporated EdU (Fig. 3B) indicating that most cells were proliferating. As the primordia were incubated for 8h with EdU, we concluded that the cycling time of these cells is comparable to the one previously measured for the LR (∼7.5h, (von Wangenheim *et al*., 2016)). To directly monitor the mitotic activity of these apically located cells before emergence, we used the cell cycle progression reporter, Cytrap (Yin *et al*., 2014), expressing the S/G2 phase marker *pHTR2::CDT1a(C3)-RFP* that we crossed to the QC25^Tg^ reporter to mark the position of the future QC cells from stage IV onward (Goh *et al*., 2016). We confirmed that most of the cells expressing the QC25^Tg^ reporter expressed the Cytrap S-phase marker indicating that they were mitotically active (Supplemental Fig. S3). We then used light sheet live imaging of developing LRP expressing *SCR::nGFP*. We observed several divisions of *SCR*-expressing apically located cells (Supplemental Fig. S3). Together, these data indicate that before emergence, apically located cells are actively proliferating and quiescence must occur once the LR emerges.

**Fig. 3.**
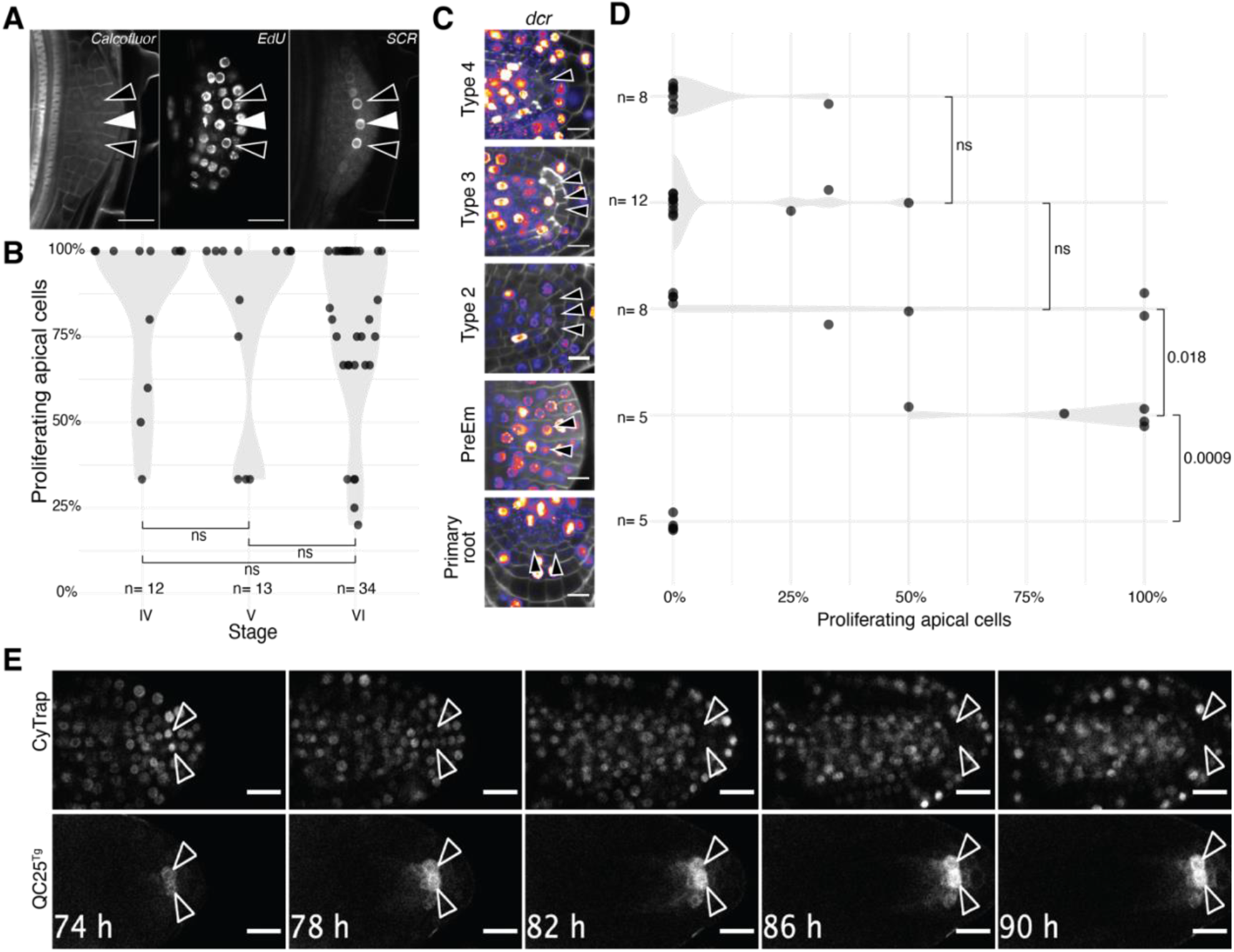
Post emergence quiescence of the LR stem cell niche. (A) Confocal sections of a pre-emergence lateral root primordium expressing *pSCR::nGFP* (*SCR*) counterstained with Calcofluor White and the S-phase label EdU. The open arrowheads indicate apically located cells expressing *SCR* and positive for EdU (8h incubation), while the filled one shows a *SCR* positive apical cell with no EdU staining. (B) Proportion of proliferating apically located cells in stage IV-VI lateral root primordia. Each point represents a primordium. The number of observed primordia is indicated. The distributions do not differ from each other according to Student’s t test with Bonferroni correction. (C) Confocal sections of primary root and lateral root apical regions at different stages of development in 10-day-old *dcr* plants counterstained with Calcofluor White (grey) and the S-phase label EdU (color). Arrowheads indicate the apically located cells. (D) Quantification of the proportion of proliferating apically located cells in the developing root apical meristem at different stages of LR development. The number of observed primordia is indicated. The level of significance of each comparison is indicated (Student’s t test with Bonferroni correction). (E) Time lapse confocal imaging of a type 2 emerged LR expressing the S-G2 phase marker CyTrap and the apical cell marker *QC25*^*Tg*^. The arrowheads track the position of two apically located cells. Note the disappearance of the S-phase marker in these cells between 74h and 82h. Hours indicate the time since induction of LR formation by gravistimulation. Scale bars: A,C: 10 µm, E: 20μm.

Emerging LR are covered by a cutin-based cuticle (Berhin *et al*., 2019) that blocks the uptake of EdU by the young LR (Supplemental Fig. S4) and consequently prevented us to analyse EdU incorporation in young emerging LR to pinpoint when post-emergence apically located cells become quiescent. To circumvent this, we monitored EdU incorporation in the *defective in cuticular ridges* (*dcr*) mutant with altered cutin biogenesis and enhanced permeability (Berhin *et al*., 2019). LR emergence in the *dcr* mutant is slower than wild type and some primordia are mis-shaped (Berhin *et al*., 2019), nevertheless as in wild type most apically located cells in *dcr* LRP incorporated EdU before emergence, indicating that the mutation does not impact the proliferative behaviour of the primordia (Supplemental Fig. S4). After emergence, the proportion of proliferating apically located cells dropped to less than 25% already in type 2 LR (<250 µm long), indicating that quiescence is quickly established after emergence (Fig. 3C). To visualise the moment when apically located cells stop proliferating, we performed live imaging of LR expressing the Cytrap/QC25^Tg^ reporters and tracked the position of apically located cells with the QC25^Tg^ marker. We observed the disappearance of the S-phase Cytrap marker in two apically located cells shortly after emergence (130µm long, Fig. 3D). Together, these results show that quiescence of the LR niche cells is established shortly after emergence.

### Biphasic acquisition of the stem cell niche organiser function in LR

The primary root stem cell niche acts as an organising centre that maintains cells in its direct contact in a stem cell state (Bennett et al., 2014; Pi et al., 2015; van den Berg et al., 1997) and generates tissues in both distal (rootward) and proximal (shootward) directions (Rahni *et al*., 2016). Whereas the position of the LR stem cell niche is set before emergence, quiescence and the final cellular layout are only established after. This raises the question of when is the niche starting to function like an organising centre able to generates tissues in both distal and proximal directions.

We first looked at the expression of markers for the proximal (enhancer trap line Q0990 (Radoeva *et al*., 2016)) or distal (*SOMBRERO*, (Bennett *et al*., 2010)) axis during LR ontogeny. In the primary root, Q0990 expression marks the vascular cells and the QC (Radoeva *et al*., 2016). In the developing LR, Q0990 is first detected at stage IV, marking the central elongated cells that will form the LR stele and is excluded from the apically located cells whose position correspond to the future stem cell niche (Fig. 4A). It is only post-emergence, in type 2 LR, that Q0990 expression starts to be visible in these apically located cells in addition to the stele, confirming that these cells only acquire a QC-like identity post-emergence (Fig. 4A). In the primary root meristem, *SOMBRERO* (*SMB*) is expressed in all the differentiated cells of the root cap and excluded from the columella stem cells (Bennett *et al*., 2010) and is thus a marker for differentiation in the distal direction. We did not detect expression of the *SMB* reporter *pSMB::nGFP* in LR before emergence. In LR that just emerged, the reporter is first detected in the distal layers of the root cap, but excluded from the cells abutting the apically located cells (as previously reported by (Du and Scheres, 2017). As LR grow longer, the number of columella layers increases (see below) and these express *SMB*, but not the cells in direct contact to the apically located cells. Taken together, it appears that proliferation in the proximal (shootward) direction is established before LR emergence, while it is only post-emergence that proliferation in the distal direction is set up (Fig. 4A). These observations suggest that it is only post emergence that columella stem cells (CSC) are specified. To verify this, we first quantified the number of columella cells layers. In the primary root four to eight layers of columella cells are homeostatically maintained through the balanced activity of the columella stem cell and the sloughing off of the outermost layer of root cap (Fendrych *et al*., 2014). Before LR emergence, we observed two layers of columella and this number progressively increased to five layers as LR matured post-emergence (Fig. 4B). We then looked at the distribution of starch granules, a marker of differentiated columella cells typically absent from the CSC (van den Berg *et al*., 1995), in developing LR. mPS-PI staining (Truernit *et al*., 2008) of LR showed that starch granules are first formed post-emergence in columella cells, confirming previous observations (Guyomarc’h et al., 2012; Kiss et al., 2002), but also revealed starch in the cells directly abutting the apically located cells. Interestingly, a layer of cells devoid of starch granules next to the apically located cells is formed only in type 5 LR (>2mm) (Fig. 4C). Together these data show that before emergence the apically located cells are able to maintain as stem cell the abutting cells in the proximal (shootward) direction but only acquire this capacity post-emergence in the distal (rootward) direction.

**Fig. 4.**
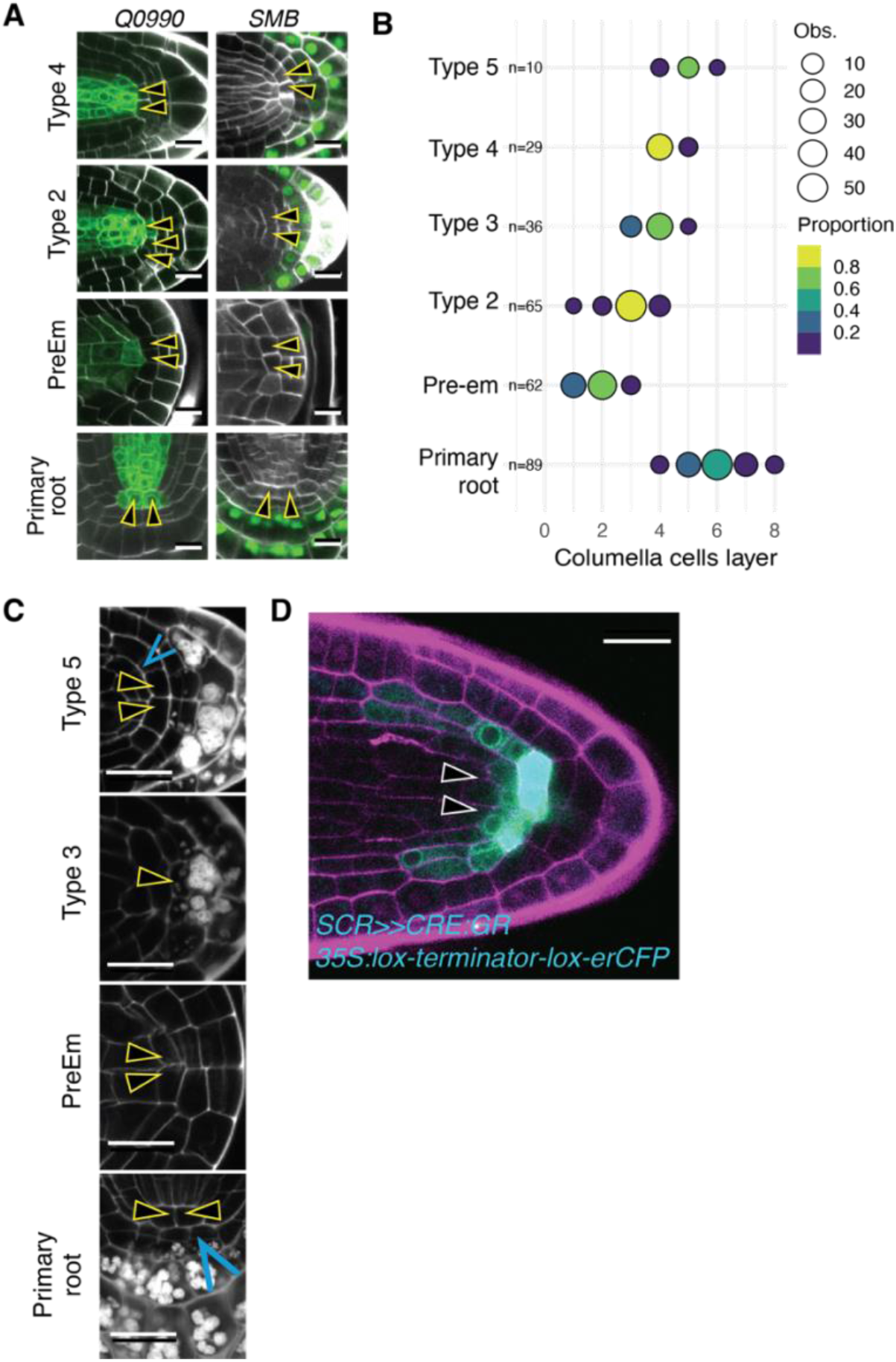
Biphasic acquisition of the stem cell niche organiser function in LR. (A) Confocal sections of calcofluor white counterstained primary root meristem and each class of LR in 10-day-old Arabidopsis expressing the proximal (*Q0990*) and distal (*SMB*) identities reporters. The open arrowheads indicate apically located cells in lateral roots and the QC in the primary root. (B) Number of columella cell layers in the primary root and for the different classes of lateral roots. The size of the circle is proportional to the number of observations (Obs.), and the colour represents the relative proportion of observations within a given class. (C) Confocal sections of mPS-PI stained primary root meristem and each class of LR in 10-day-old Arabidopsis. The open arrowheads indicate apically located cells in lateral roots and the QC in the primary root while the blue arrow point the layer of columella stem cells. (D) Confocal section of a PI counterstained type 2 lateral root (length: 112µm) 38h after generation of clones in the *SCR* domain by dexamethasone treatment. Arrowheads indicate provascular initials resulting from the division of the *SCR*-expressing apically located cells. Scale bars are 10µm (A, D) and 20µm (C).

To confirm this, we induced clones in the *SCR* domain and checked whether the clones derived from the central cells marked the proximal and/or distal domains. Observation of post-emergence LR expressing the *p35S::lox-Ter-lox:erCFP / pSCR::CRE-GR* construct (Efroni *et al*., 2016) and treated with DEX for 38h resulted in erCFP signal in the apically located cells, the CEIs, and some endodermal cells. We also observed erCFP expression in the provascular cells directly adjacent to the apically located cells but none in the cells located distally (Fig. 4D). Thus, the LR stem cell niche acquires its ability to bidirectionally generate tissues in two phases: it contributes to growth in the proximal direction before emergence and in the distal direction only post-emergence, concomitantly to the establishment of its mitotic quiescence and expression of specific marker genes.

## Discussion

Here, we characterised the ontogeny of the root apical meristem (RAM) in lateral root, with a particular emphasis on the QC. We used reporters for tissue identity, live imaging, lineage tracing and cell proliferation assay and conclude that while the QC position and its role in proximal (shootward) growth are set early during LR formation, its molecular identity, cellular organization and quiescence are only fixed after LR emergence. The different attributes of the primary RAM QC are thus progressively acquired during LR formation, suggesting a progressive maturation process of this post-embryonic meristem.

Despite the similarities in cellular organisation and gene expression profiles between the primary and LR RAM and the essential role played by the QC in organising and coordinating the primary RAM (Rahni *et al*., 2016), there were no comprehensive analysis of the ontogeny of the QC in LR. Previous reports analysed either the expression patterns of genes of the primary RAM during LR formation (Tian et al., 2014a) or focused on cell differentiation associated with acquisition of gravitropic and phototropic responses in LR (Guyomarc’h et al., 2012; Kiss et al., 2002; Waidmann et al., 2020). Our analysis reveals interesting insights into the mechanisms of LR RAM maturation.

Like in the primary RAM, the pattern of auxin signalling and of *SCR* and *WOX5* expression defines the position of the QC during the early stages of LR development. This result agrees with a prior report (Goh *et al*., 2016) and supports the view that a key transition in LR organogenesis occurs at the stage IV-V transition, when the primordium breaks through the overlying endodermis layer, becomes radially symmetric (Lucas *et al*., 2013) and acquires autonomy relative to the auxin fluxes from the parent root (Laskowski *et al*., 1995). Yet, our observations nuance the conclusion that the LR QC is established at this early stage (Goh *et al*., 2016). Indeed, we observe variation in the cellular organisation of the LR RAM prior to emergence and a primary root like cellular organisation of the RAM is only fixed once the LR has emerged. Our analysis of cell proliferation using EdU incorporation or long term imaging revealed that quiescence of the apically located cells in the LR RAM is only established after emergence. This quiescence is concurrent to the onset of expression of most molecular markers associated with the primary RAM (*AGL42, BRAVO*, QC184 and QC25^ET^). An exception being QC25^Tg^ which marks the position of the apically located cells from stage IV/V albeit a stage when these cells still proliferate. The difference of behaviour between QC25^Tg^ and QC25^ET^ reporters suggests that whereas the positional cues responsible for the localised expression of the QC25^Tg^ reporter are present in its promoter sequence, they are overridden in the enhancer trap line until the cells become quiescent. In addition to revealing the existence of genomic regulations associated with the quiescence of the apically located cells, this results support the temporal decoupling between positioning the prospective LR QC cells and their actual maturation and entry into quiescence. Our results support that the position of the LR QC is set before emergence but only activated post-emergence as it had been suggested earlier (Celenza *et al*., 1995; Malamy and Benfey, 1997). A recent single cell analysis of the developing LR has shown that expression of genes typically associated with the QC in the primary RAM can be readily detected in stages I to III LR, yet distributed amongst different clusters that only form a distinct coherent population at later stages (Serrano-Ron *et al*., 2021). Our observation of *WOX5-YFP* expression in most cells of stage II LR supports this notion of a progressive specification of cell identities in the early phases of LR organogenesis. This early expression of *WOX5* is raising questions about of the role played by this essential factor for QC identity at early stages. Interestingly, (Goh *et al*., 2016) reported a very specific onset of WOX5 expression in four apically located cells at stage IV/V, in apparent contradiction to our and others observations (Ditengou *et al*., 2008; Du and Scheres, 2017). The absence of essential cis-regulatory element(s) in the transcriptional reporter used by Goh et al. (*pWOX5::n3GFP*) compared to the YFP-tagged genomic clone used (Pi *et al*., 2015) could account for the differences in expression pattern.

Current models assign to the QC an organizer function that signals to the surrounding cells and actively maintains their “stemness”. An alternative view is that the position of the stem cells organizer results from two opposing axis that reciprocally control each other’s differentiation (Rahni *et al*., 2016). In this so called “stagnation” model, the QC corresponds to the location where cells are not displaced by their neighbours as result of the activity of shootward and rootward gradients of division and differentiation. Where these two gradients meet, cell division is minimal and a stagnation zone is created on either side of which more rapidly dividing cells are bound to displaced their daughter cells in opposite directions (Rahni *et al*., 2016). In this model, the stemness of stem cells is the result of a passive mechanical process and does not rely on quiescence of the QC. Our results show that the growth axis of the LR are established in distinct phases with the proximal (shootward) growth axis established pre-emergence and the distal (rootward) one after emergence, concomitantly to the stabilisation of number of prospective QC cells, the slowdown of their cell division rate and the activation of QC specific markers. These data support the stagnation model in which the position and maturation of the LR QC cells is fixed once the two opposite gradients are established. Activation of QC specific identities and entry in quiescence would be an emerging property of the slowdown in cell division rate at the position where the two growth axis meet. Quiescence, a late emerging property in LR, would thus not be instrumental in organizing the stem cell niche, but a results of the two diverging growth axis, alike the region of mostly calm weather at the center of a cyclone results from diverging convection flows (Rahni *et al*., 2016). This shift in the importance of the QC slow mitotic activity is supported by the observation that loss of QC-quiescence has no effect on root growth (Cruz-Ramírez *et al*., 2013).

## Material and Methods

### Plant material and growth conditions

We used the previously described lines: *DR5v2::3xYFP:NLS* (sC111) (Vilches Barro *et al*., 2019), *pTCSn::2xVenus:NL*S (El Arbi *et al*., 2021), *gWOX5::WOX5:YFP* in *wox5-1* (Pi *et al*., 2015), *pAGL42::truncAGL42:GFP* (Nawy *et al*., 2005), *pBRAVO::BRAVO:GFP* (Vilarrasa-Blasi *et al*., 2014), *pQC184::GUS* (in the gWOX5::WOX5:YFP in *wox5-1*) (Pi *et al*., 2015), *pQC25er::CFP* (QC25^Tg^) (Goh *et al*., 2016), Cytrap (*pHTR2::CDT1a (C3)-RFP, pCYCB1::CYCB1;1-GFP*) (Yin *et al*., 2014) and *pSCR>>CRE:GR/35S:lox-terminator-lox-CFP* (Efroni *et al*., 2016) The enhancer trap lines Q0990 (Radoeva *et al*., 2016), QC25^ET^ (Sabatini *et al*., 2003) were described previously. For crosses between lines, F1 generation was used. The *dcr* mutant was previously described (Berhin *et al*., 2019). Seeds were surface sterilized (Sodium hypochlorite 5 % and 0.05 % Tween 20 or Ethanol 70 % and 0.05% Triton X-100) and placed on ½ Murashige and Skoog (MS) medium containing 0.8 % agar (Duchefa). Following stratification (4°C in the dark, >t 24 h), seedlings were grown at 22°C vertically under long day conditions (16 h light / 8 h dark). For the lineage tracking experiments 15µM dexamethasone were used for 24h.

### Construction of vectors and plant transformation

The transcriptional reporters *pSCR::nGFP (pSCR::H2B-3xGFP::tRBCS)* and *pSMB::nGFP* (*pSMB::H2B-3xGFP::tRBCS)* reporters were generated using GreenGate assembly (Lampropoulos *et al*., 2013). For *pSCR*, 2114bp upstream of ATG were used and 3246bp for *pSMB. Agrobacterium tumefaciens* (Agl-0) based plant transformation was carried out using the floral dip method (Clough and Bent, 1998). All plant lines examined were homozygous if not indicated otherwise. Homozygosity was determined by antibiotic resistance and verification of the fluorescent fusion proteins at the microscope.

### Microscopy

Unless otherwise specified, imaging was performed on seedlings fixed for 30 min with PFA 4% in PBS 1X at room temperature, Calco Fluor white staining was performed as previously described (Ursache *et al*., 2018). Seedlings were cleared with either 50% glycerol for 24h or ClearSee solution for up to a week (Kurihara *et al*., 2015). Root meristems were imaged on medial sections on a Leica SP8 confocal microscope with 63x, NA = 1.4 oil immersion objective or a 40x, NA = 1.3 oil immersion objectives. Calcofluor White fluorescence was detected using the 405nm excitation laser line, and 425-475nm emission range. The number of apically located cells is obtained by counting all cells between the cortex-endodermis initial. For Edu incorporation, Alexa fluor 647nm detection was achieved using the 638 nm excitation laser line in the 650-695nm range. For the Cytrap cell cycle progression reporter (Yin *et al*., 2014), the S and G2 phases marker pHTR2::CDTa (C3)-RFP was detected using the 552nm nm excitation laser line in the 600nm to 650nm emission range. The G2/mitosis phases marker was detected using the 488nm excitation laser line in the 500 to 550nm emission range. For live imaging, a Leica SP8 was used with a 20X NA= 1.2 oil immersion objective. Lateral root growth was induced by 52h of gravistimulation (180°) in seedlings of 4 to 6 days after germination (dpg). Seedlings were placed in an imaging chamber and covered with a piece of ½ MS medium for imaging, as described in (Marhavy and Benkova, 2015). Stacks were acquired every 2 hours with 1µm z-steps. For live imaging of the *SCR* reporter, the *pSCR::nGFP* reporter was crossed to a line expressing *pUBQ10::PIP1,4:GFPx3; pGATA23::H2B:nCherryx3* (stPVB003, (Vilches Barro *et al*., 2019) and imaging was done on a MuViSPIM (Luxendo, Bruker, Germany) light sheet microscope equipped with a Nikon NIR Apo 40X NA=1.2 water objective for detection. Movies were acquired with a z-step of 0.250µm, and a temporal resolution of 30 min.

### Histochemical analysis

Staining of starch granule in the columella cells was performed as described previously (Truernit *et al*., 2008). GUS staining was performed as described previously (Maizel and Weigel, 2004) followed by fixation in 4%HCl, 20% methanol solution, for 15 min at 70°C, followed by 7%NaOH and 60% ethanol for 15 min at room temperature. Seedlings were then cleared in successive ethanol baths for 10 mins (40%, 20%, 10%), followed by 10 min incubation in 25% glycerol and 5% ethanol. Finally seedlings are placed in 50% glycerol and stored at 4°C until imaging. Seedlings were mounted in 50% glycerol for imaging with DIC microscopy using an Axio Imager. M1 (Carl Zeiss, Oberkochen, Germany) with a 40X objective.

### EdU incorporation

For Edu incorporation in LRP before emergence, LR formation was induced by 30h, 36h, or 42h of gravistimulation, and seedling were transferred on plates containing 1µM of EdU in ½ MS plates for the last 8h of the gravistimulation period. For Edu incorporation in emerged lateral root (in *dcr* mutant), LR was induced by 48h, 72h, 96h or 120h of gravistimulation, and EdU incorporation was performed during the last 24h of the gravistimulation using 0.5 µM of EdU in ½ MS plates. EdU detection was performed using a slightly modified version of the previously described protocol (Kotogány *et al*., 2010). Briefly, seedlings were fixed for 15 min at room temperature in the dark with 4% PFA in PBS, washed 2 times 1 min with 3% BSA in PBS, incubated 20 min with 0.5% Triton X-100 in PBS and washed 2 times 1 min with 3% BSA in PBS, then incubated with the Click-iT™ (Click-iT™ EdU Cell Proliferation Kit for Imaging, Alexa Fluor™ 647™ Invitrogen, Thermofisher, C10340) reaction cocktail for 30 min at room temperature in the dark, rinsed with PBS and washed with 3% BSA in PBS 3 times for 1 min. Seedlings were then washed 2 times with PBS and cell walls were stained with 0.1% Calco fluor white (Fluorescent brightener 28 F3543-1G Sigma-Aldrich) in PBS for 30 min in the dark. After rinsing, seedlings were transferred to 50% glycerol for at least 16h before imaging. Apically located cells showing EdU incorporation or expressing the pHTR2::CDTa (C3)-RFP marker (Cytrap) were considered as proliferative. The proportion of proliferating apically located cells was obtained by dividing the number of proliferative apically located cells divided by the total number of apical cell on that section.

## Supplementary data

Supplementary data are available online.

Fig. S1. Expression of WOX5 in early lateral root primordia

Fig. S2. Expression of SCR during germination

Fig. S3. Apically located cells stop to proliferate post-emergence

Fig. S4. EdU incorporation and cell proliferation in LR of wild type and dcr mutant

## Acknowledgements

We thank J. Dubrovsky for critically reading the manuscript, and the Caño-Delgado, Efroni, Greb, Laplaze, Laux, Lohmann, Nawrath and Palatnik labs for sharing resources. The authors gratefully acknowledge the data storage service SDS@hd supported by the Ministry of Science, Research and the Arts Baden-Württemberg (MWK) and the German Research Foundation (DFG) through grant INST 35/1314-1 FUGG and INST 35/1503-1 FUGG.

## Author contribution

Conceptualization: AM

Data Acquisition and curation: BB, AM

Formal Analysis: BB, AM

Funding Acquisition: AM

Methodology: AM, BB, LB, ML

Project Administration: AM

Resources: BB, LB, ML

Supervision: AM

Visualization: BB, AM

Writing – Original Draft: BB, AM

Writing – Review and Editing: BB, AM

## Conflicts of interests

The authors declare no competing interests.

## Funding

This work was supported by the DFG FOR2581.

## Data availability

The data supporting the findings of this study are available from the corresponding author upon request.

## Supplemental material for

**Fig. S1.**
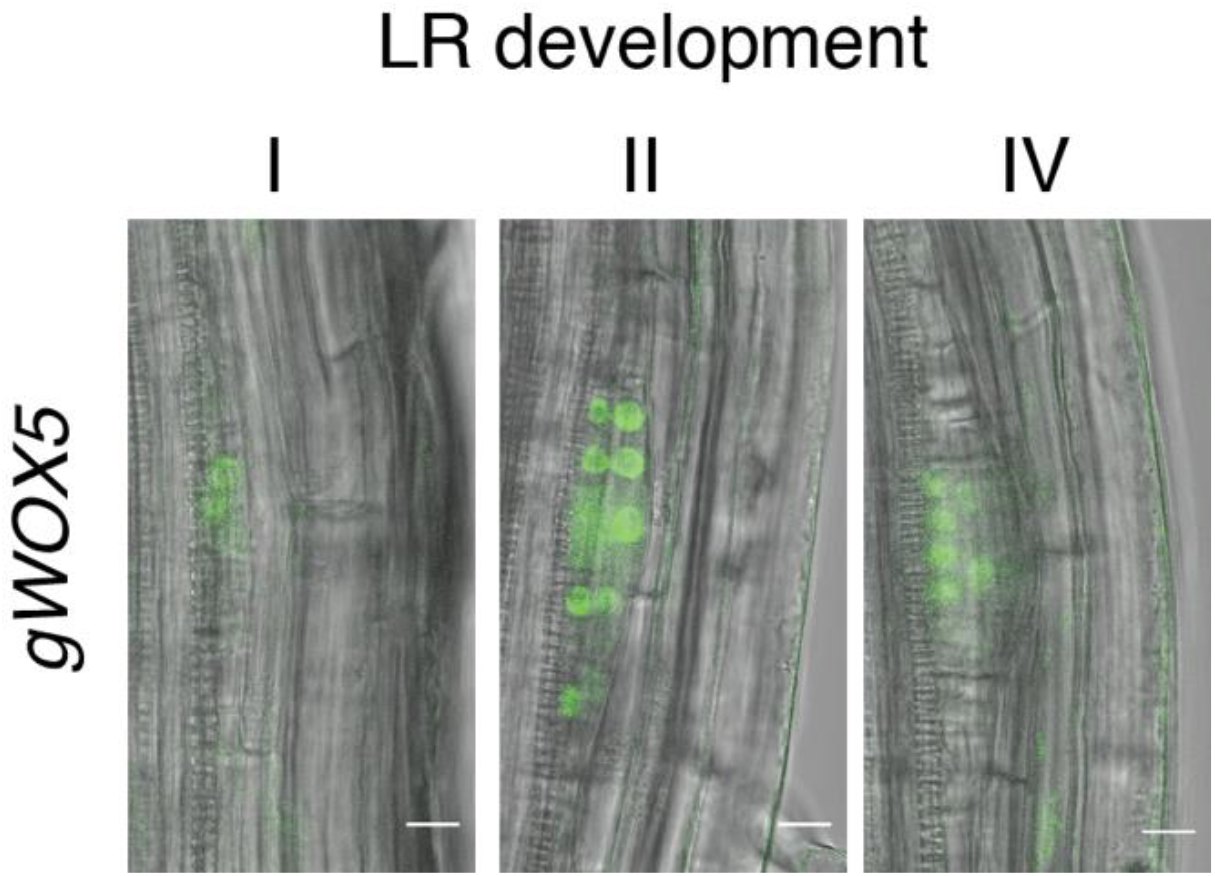
Expression of *WOX5* in early lateral root primordia. Confocal section of 10-day-old *gWOX5* at stage I, II and IV of lateral root development. Scale bar 10µm.

**Fig. S2.**
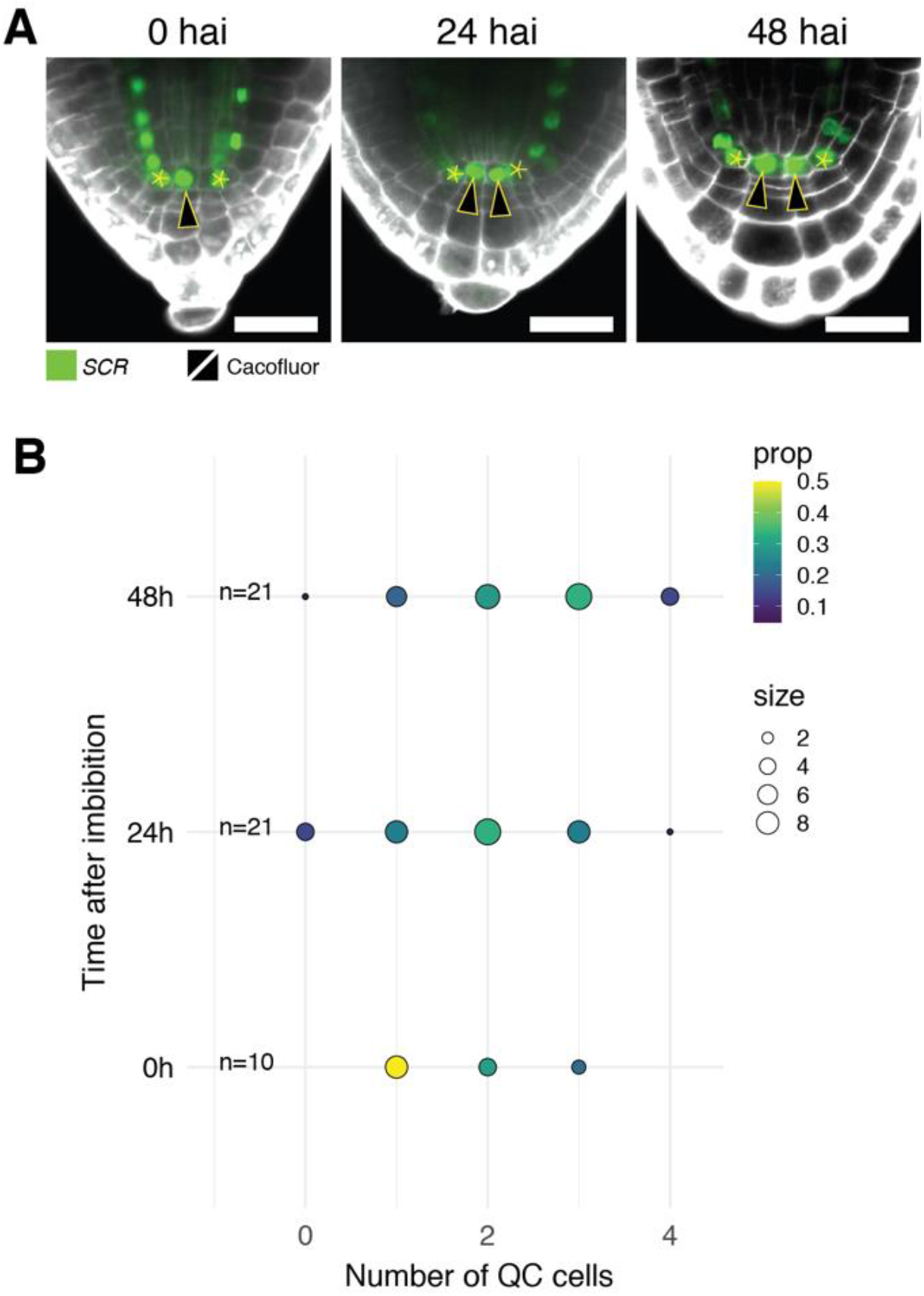
Expression of *SCR* during germination. (A) Confocal section *SCR* (*pSCR::nGFP*) embryos at 0, 24 and 48h post seed imbibition. The QC cells (arrowheads) are indicated. Scale bar 10µm. (B) Number of QC at 0, 24 and 48h post seed imbibition. The size of the circle is proportional to the number of observations (Obs.), and the colour represents the relative proportion of observations within a given class. Total number of observations is indicated.

**Fig. S3.**
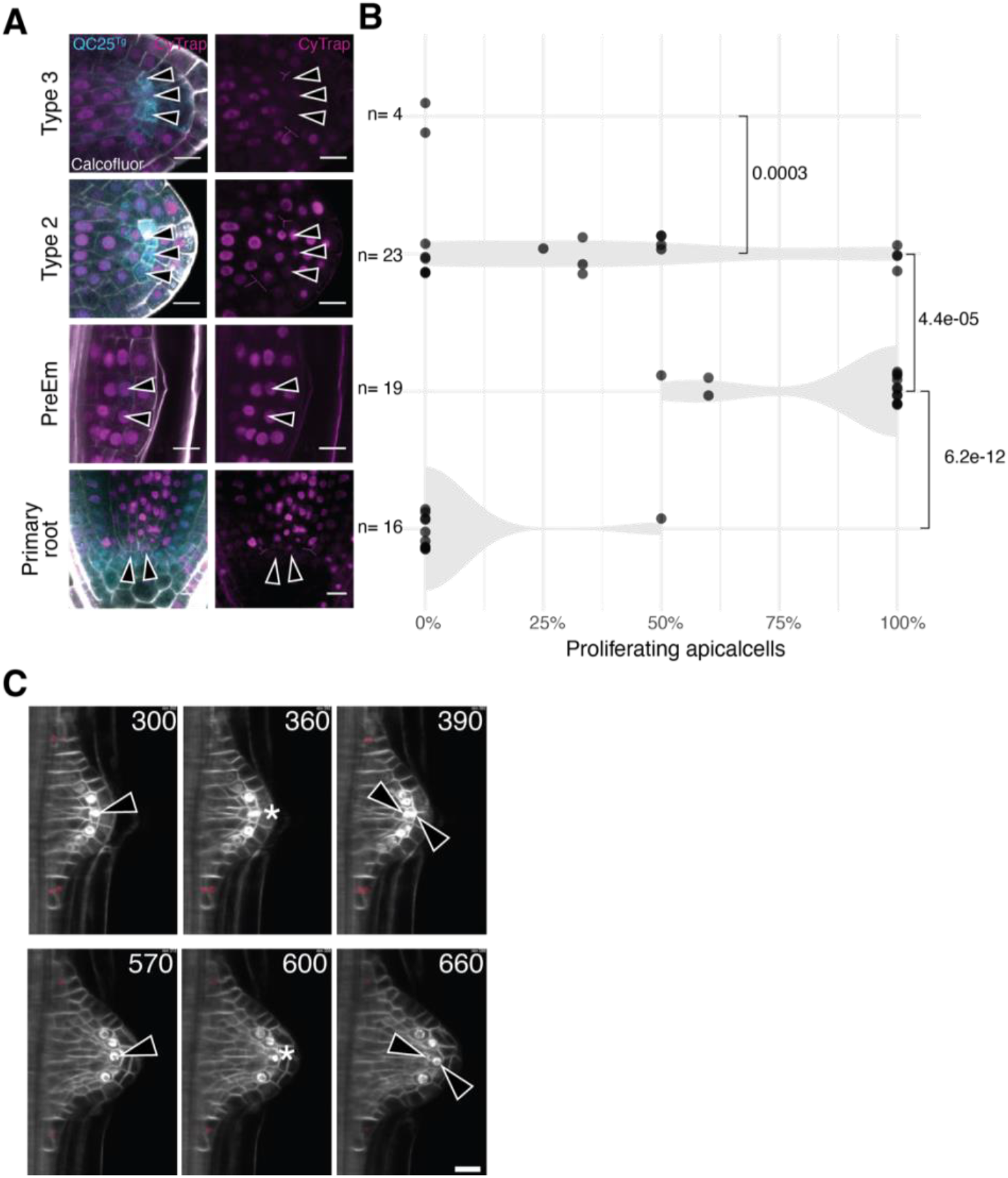
Apically located cells stop to proliferate post-emergence. (A) Confocal sections of primary root and lateral root apical regions at different stages of development in lines expressing the S-phase marker CyTrap, the apical cell marker QC25^Tg^ and counterstained with Calcofluor White. Arrowheads indicate the apically located cells. (B) Quantification of the proportion of proliferating apically located cells in the primary root meristem and different stages of LR development. The total number of observations and the statistical significance (Wilcoxon test) of the difference between groups are indicated. (C) Time lapse light sheet imaging of a developing LR expressing *SCR* (*pSCR::nGFP*) and a plasma membrane marker (PIP1;4-GFP, see methods). Arrowheads point at apically located cells, while asterisks mark mitosis. Two divisions are observed 360 and 600 minutes after the start of the recording. Scale bars: 10μm.

**Fig. S4.**
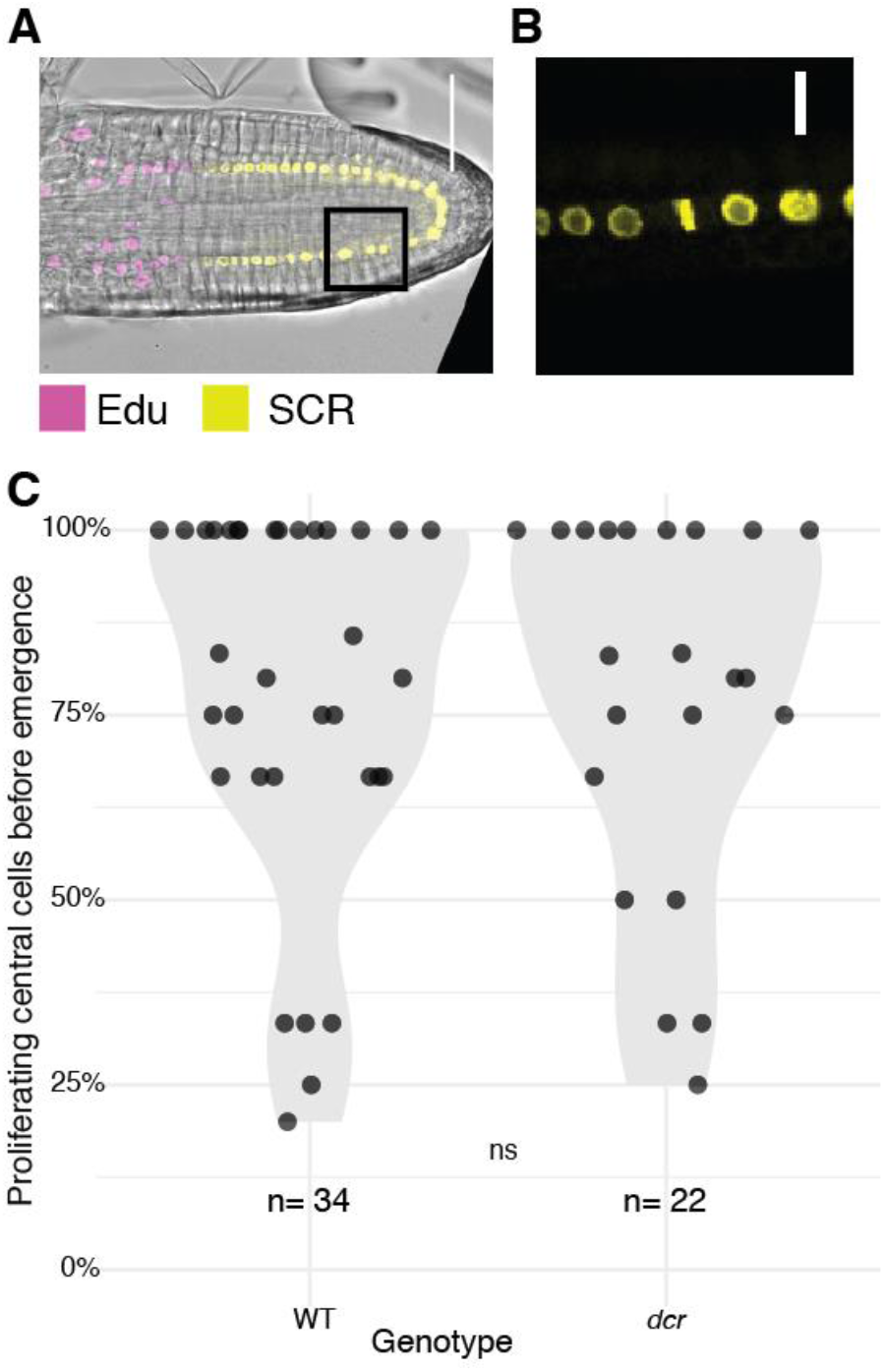
Lack of EdU incorporation in young wild type emerged lateral roots and comparable proliferation of the lateral roots apically located cells in wild type and *dcr*. (A) Confocal section of wild type 10-day-old *SCR* (*pSCR::nGFP*) emerged lateral root counterstained with EdU. (B) Close up on the region boxed in (A) showing cell division despite lack of EdU incorporation. (C) Proportion of proliferating apically located cells in stage IV-VI LRP in wild type (WT) and *dcr* mutant. Each point represents a primordium. The distributions do not differ from each other according to Wilcoxon’s test.

## Notes

### Competing Interest Statement

The authors have declared no competing interest.

